# Elephants inhabiting two forested sites in western Uganda exhibit contrasting patterns of species identity, density, and history of hybridization

**DOI:** 10.1101/2025.05.13.653790

**Authors:** Claire K. Goodfellow, Daniella E. Chusyd, Savanah R. Bird, Dennis Babaasa, Colin A. Chapman, Jena R. Hickey, Mireille B. Johnson, Charles Kayijamahe, Richard Mutegeki, Patrick A. Omeja, Charles Tumwesigye, Eustrate Uzabaho, Samuel K. Wasser, Michael D. Wasserman, Caitlin P. Wells, Nelson Ting

## Abstract

Elephant populations across much of Africa face severe rates of decline due to poaching and habitat loss. The recent decision by the International Union for the Conservation of Nature (IUCN) to separately list African forest (*Loxodonta cyclotis*) and savanna (*L. africana*) elephants on the IUCN Red List both highlights the different threats of extinction faced by these two species and emphasizes the need for genetic data to classify taxonomically undefined populations across the continent. This includes western Uganda – a region that harbors the largest known modern hybrid zone between the two species. We combined a new high-throughput amplicon sequencing (HTAS) approach with fecal DNA-based Capture Mark Recapture (CMR) analysis to infer the population sizes and species compositions of elephants living in two forests. We demonstrate that Kibale National Park hosts a relatively large elephant population (573 individuals, 95% CI: 410 to 916; 0.72 elephants/km^2^) composed primarily of hybrids (81.5%) and savanna elephants (17.7%), while Bwindi Impenetrable National Park hosts a smaller population (96 individuals, 95% CI: 64 to 145; 0.29 elephants/km^2^) composed of forest elephants (86.8%) and hybrids (13.2%). We then sequenced maternally inherited (mtDNA) and paternally inherited (AMELY) genetic markers and found that the two parks’ populations exhibit different patterns of sex-linked genetic variation. The contrasting patterns of species identity and genetic variation between these parks demonstrate different histories of hybridization and highlight the importance of site-specific monitoring where elephants are taxonomically undefined.

## Introduction

Recent advances in sequencing technology have revealed a surprisingly high degree of both ancient and ongoing hybridization across a wide range of taxa (Taylor & Larson, 2019). This interspecific gene flow can influence the adaptation, evolution, and survival of species, and can be a natural evolutionary process (Arnold, 1992) and/or a consequence of anthropogenic activity (Grabenstein & Taylor, 2018). For management, the increased ability to detect these hybrids using genetic tools demands the use of increasingly complex decision-making frameworks by conservation practitioners (Allendorf et al., 2001; Jackiw et al., 2015). Small populations of rare or endangered species are especially prone to genetic swamping from hybridization with closely related domestic, introduced, or more abundant sympatric taxa (Quilodrán et al., 2020). Additionally, because many wildlife regulations are designated at the species level, hybridization can compromise legal protections for the hybrid offspring of endangered species (Allendorf et al., 2001).

Conservation authorities face this challenge of hybridization in protecting Africa’s elephants, which are declining across the continent due to poaching, human-wildlife conflict, and habitat loss (Thouless et al., 2016). Both morphological (Grubb et al., 2000) and genetic (Roca et al., 2001; Palkopoulou et al., 2018) data support the recognition of two African elephant species: the forest (*Loxodonta cyclotis*) and savanna elephant (*Loxodonta africana*). These two distinct lineages diverged between 2.6 and 5.6 million years ago (Palkopoulou et al., 2018) and differ in distribution (Mondol et al., 2015; Kuhner et al., 2025), behavior (Athira & Vidya, 2021), and life history traits (Turkalo et al., 2018). Yet, due in part to uncertainty surrounding both the prevalence of hybridization and how hybrids might be affected by a two-species classification, most conservation, government, and regulatory agencies were hesitant to formally acknowledge separate species. This is because “lumping” African elephants into one conservation unit simplified both international law enforcement efforts and local management decisions and prioritized full protection for hybrids. However, it also obscured the population numbers for each species, which can lead to negative conservation outcomes (Morrison et al., 2009).

In 2021, the IUCN African Elephant Specialist Group officially split its listing status for Africa’s elephants, which were previously listed together as “Vulnerable” (Hart et al., 2021). This change required separate population assessments for the two species and resulted in a “Critically Endangered” status for forest elephants (Gobush et al., 2021b), and an “Endangered” listing status for savanna elephants (Gobush et al., 2021a). While this decision highlights the different threats of extinction faced by the two species, it also poses a series of new challenges (Hart et al., 2021), including a crucial need to generate high-resolution, species-level distribution data for taxonomically undefined priority sites across the African continent (Bauer et al., 2021).

Western Uganda contains several priority sites for elephant conservation embedded within the largest known modern African elephant hybrid zone (Mondol et al., 2015). While elephants ranged freely throughout this region as recently as 1929 (Brooks & Buss, 1962), they are now largely confined to a network of protected areas embedded in a highly populated agricultural matrix (UWA, 2016). The dense vegetation and steep topography of several of these protected areas have made traditional sight-based elephant census methods unfeasible; hence reliable census counts and species distribution data are lacking (Plumptre et al., 2008). The line-transect distance sampling (LTDS) method is typically used in forested areas, relying on both accurate dung counts along a series of replicated transects and seasonally accurate defecation and dung decay rates for every surveyed site (Barnes, 1996). While this method is well established and widely used, it does not identify elephants to species, making it less useful in taxonomically mixed or undefined populations. One alternative that enables inference of both census population size and species distribution is a fecal DNA-based Capture Mark Recapture (CMR) approach (Hedges et al., 2013; Brand et al., 2020; Laguardia et al., 2021b). Like the LTDS method, this CMR approach relies solely on elephant dung. However, it does not require previous knowledge about site-specific parameters such as defecation and dung decay rate to accurately infer population size and thus shows promise as a method for dual inference of species level genetic variation and abundance.

Here, we implement a fecal DNA-based CMR approach to update population estimates for elephants in two high conservation priority forested sites in Western Uganda. Specifically, we apply a high-throughput amplicon sequencing approach to infer the population sizes, species compositions (i.e., forest, savanna, hybrid), and patterns of nuclear and sex-linked genetic variation for elephants at these sites. The goal was to establish accurate baseline data on how many elephants exist in these parks and to provide clarity on how these populations should be classified on national and international species inventories.

## Study Area

This study was conducted across Kibale National Park (KNP) and Bwindi Impenetrable National Park (BINP), two forested protected areas in Western Uganda. Both sites are in the Greater Virunga Landscape (Fig. 1), a globally important region for its high rates of endemism and biodiversity. KNP is a 795-km^2^ protected area near the foothills of the Rwenzori mountains (0°13’-0°41’N and 30°19’- 30°32’ E). Its elevation ranges from 1,110 to 1,590 m above sea level, and its vegetation is classified as medium-altitude semi-deciduous forest, medium-altitude moist evergreen forest and grasslands (Langdale-Brown et al., 1964). BINP is a 331-km^2^ protected area located in the Kigezi highlands (0°53’-1°08’ S; 29°35’-29°50’ E). Its elevation ranges from 1,160 to 2,600 m above sea level, and its rugged terrain is characterized by steep-sided ridges and narrow, poorly-drained valleys (Butynski & Kalina, 1993). The vegetation is classified as medium-altitude moist evergreen forest and high-altitude submontane forest (Langdale-Brown et al., 1964). Annually, the region experiences wet seasons from March – May and September – November, and dry seasons from December – February and June – August (Plumptre et al., 2008).

**Figure 1.**
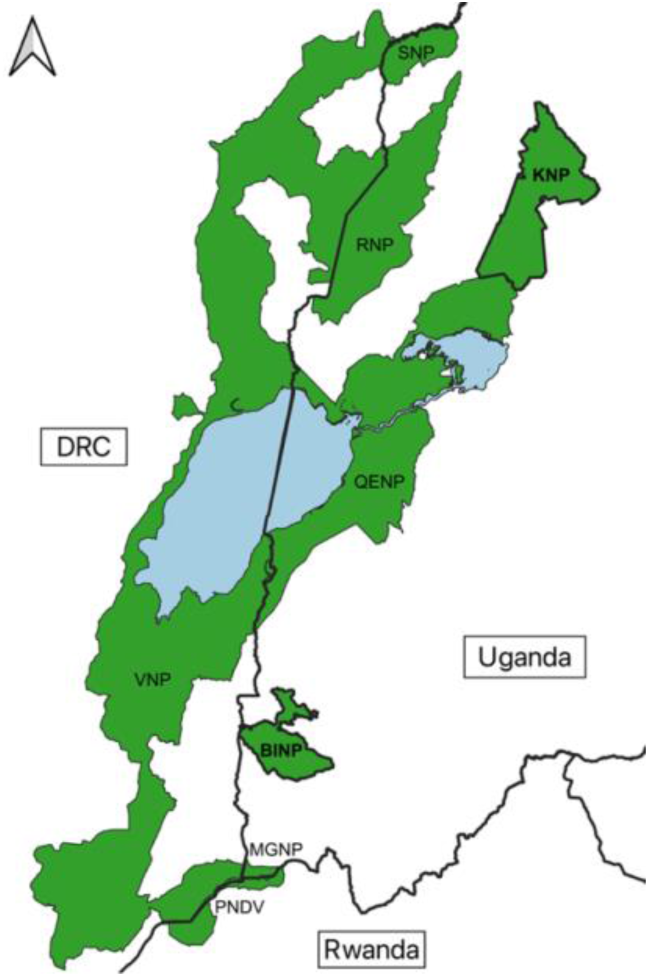
Map showing protected areas which together make up the Greater Virunga Landscape. This network spans the borders of Uganda, Rwanda, and the Democratic Republic of the Congo (DRC) and includes both Kibale National Park (KNP) and Bwindi Impenetrable National Park (BINP), two forested protected areas located in Western Uganda. Abbreviations for the parks above are: KNP - Kibale National Park; BINP - Bwindi Impenetrable National Park; MGNP - Mgahinga Gorilla National Park; RNP - Rwenzori National Park; VNP - Virunga National Park; PNDV - Parc National des Volcans; SNP - Semuliki National Park; QENP - Queen Elizabeth National Park. (Map generated in QGIS v.3.30. Data Source: National Forest Authority, 2007).

## Methods

### Sampling Scheme

Dung samples were non-invasively collected from both parks between 2018-2021, with a total of nine months of collection in KNP and six months in BINP. The sampling strategy was systematic and based on weekly rotation through different cells and/or hotspots in a spatial grid overlaid on each park, although the details of each strategy differed slightly by year/season due to differences in available personnel and/or COVID-19 restrictions. Only samples judged to be less than 48 hours old by appearance were collected. All samples were stored in RNAlater at ambient temperature before being moved to a −20°C freezer within six weeks of collection. See the Supplemental Methods for a detailed description of the sampling scheme and methods for each park.

### Genotyping

Genetic lab work and analyses were conducted in the Ting Lab at the University of Oregon. DNA was extracted using the QIAamp Fast DNA Stool Mini Kit (Qiagen) with slight modifications to the manufacturer protocol. Each DNA extract was molecularly sexed following Ahlering et al. (2011). To examine sex-linked patterns of genetic variation and infer patterns of hybridization, we also sequenced two uniparentally-inherited genetic markers: a 319 bp region of the mitochondrial control region (Brandt et al., 2012) for all individuals and a 719 bp region of the Y-linked, intronic ameloglobin (AMELY) gene (Mondol et al., 2015) for all males. Finally, all samples were genotyped at 14 biparentally inherited microsatellite loci previously shown to be polymorphic in African elephants. Genotyping was conducted in triplicate for each sample using a two-step, high-throughput amplicon sequencing (HTAS) approach (Barbian et al., 2018). Sequencing of multiplex pools in dual-indexed libraries was spread across four Illumina MiSeq runs at the University of Oregon Genomics & Cell Characterization Core Facility (GC3F) using v3 chemistry at 375 forward read cycles. See Supplemental Methods for a detailed protocol for library preparation and sequencing.

Illumina libraries were demultiplexed by sample, and adapters were trimmed and quality filtered (Q>20) using cutadapt (Martin, 2011). Per-locus demultiplexing and genotype calling were then implemented in CHIIMP (Barbian et al., 2018; Supplemental Methods). To validate genotype calls, we generated histograms of read lengths for each locus using a custom python script and manually called genotypes based on size. We required that consensus genotypes be confirmed in two or more replicates for heterozygous calls, and in three replicates for homozygous calls. GenAlEx v. 6.51b2 was used to generate standard metrics of genetic diversity (e.g., H_o_, H_e_) and differentiation (F_ST_), to test for deviations from Hardy Weinberg Equilibrium (HWE), and to identify unique individuals. P(ID) and P(ID)_sib_ – the probability that two individuals or siblings, respectively, randomly drawn from a population will have identical genotypes across multiple loci – were calculated for our panel of 14 loci using GenAlEx.

### Census Population Size Estimates

We used the R package *capwire* v. 1.3 to implement CMR analyses for each park. The statistical framework implemented in this package is optimized for use with non-invasive genetic samples and generates population size point estimates based on capture class frequency data (Pennell et al., 2013). To avoid pseudo-replication, we collapsed repeat detections of the same individual in the same sampling occasion to single detections. We then fit two different models to our data for each park: the Equal Capture Model (ECM) and the Two-Innate Rates Model (TIRM), but we only report on results from the ECM as it is the simpler model and likelihood ratio tests (N=1,000 bootstraps) determined no differences between the two. We generated point estimates for population size for each of the two parks using a maximum population size of 3,000 individuals and implemented parametric bootstrapping (N=1,000) to generate 95% confidence intervals around each point estimate.

### mtDNA and Y chromosome Clade Assignment

Sequences for the mitochondrial control region and Y-linked (AMELY) gene were trimmed, assembled and aligned using Geneious v.2023.1 (Kearse et al., 2012). Each mtDNA sequence was classified as either “S-clade” or “F-clade” based on a set of species-diagnostic SNPs representing deeply diverged (∼5.5 million year) maternal clades from the savanna and forest elephant lineages, respectively (Ishida et al., 2013). Unique mitochondrial haplotypes were assigned haplotype names based on previous detections archived in the *Loxodonta Localizer* database (Zhao et al., 2019). AMELY gene sequences were similarly classified as either “forest- typical” or “savanna-typical” based on species-diagnostic SNPs previously used to examine paternal inheritance in African elephants (Mondol et al., 2015).

### Species Classification

A set of 19 putatively pure samples were genotyped and analyzed alongside the samples from KNP and BINP to serve as positive controls for species assignment analyses (*L. cyclotis*, n=9; *L. africana*, n=10; Table S1). Bayesian clustering analysis (K=1-5) was implemented in STRUCTURE v. 2.3.4 (Pritchard et al., 2000) using the Admixture model. Analyses were run at two different scales: 1) for each park separately; and 2) for the two parks combined. At each scale, we also implemented analyses with and without the LOCPRIOR parameter set to TRUE. For all analyses, the parameters INFERALPHA and POPALPHAS were set to TRUE. Five MCMC replicates were run per K, each with an initial burn-in of 200,000 followed by 4,000,000 iterations.

Following STRUCTURE analyses, we identified the K with the highest support using STRUCTURE HARVESTER v. 0.6.94 (Earl & vonHoldt, 2012), and classified individuals as pure forest, pure savanna, or hybrid using two different methods. For the first method, we implemented a standardized cutoff to classify individuals based on STRUCTURE output: individuals with a posterior ancestry proportion Q ≥ 0.9 for either of the two putative populations (K=2) were classified as a “pure” species for that cluster. This cutoff has been demonstrated to provide an optimal threshold for distinguishing between hybrid and parental types when using a small number of molecular markers (van Wyk et al., 2017). Individuals with Q < 0.9 for either ancestry-specific cluster were considered admixed and classified as hybrid. For our second classification method, we used the R package EBHybrids v. 0.991 (Mondol et al., 2015) to classify individuals by species and assign each sample a hybrid probability. This package uses an empirical Bayes method to quantify the likelihood of a sample being from a pure savanna elephant, pure forest elephant or hybrid based on STRUCTURE results, locus-specific allelic dropout rates, and a reference panel of putatively un-admixed individuals. As has been previously done (Bonnald et al., 2021), each individual was classified into a species category at three different likelihood thresholds (50%, 80%, and 95%).

## Results

### Sample Collection and Individual Identification

A total of 399 samples from Western Uganda (KNP, n=256; BINP, n=143) were collected and analyzed. The cumulative P(ID) for the 14 loci used was 1.2 x 10^-9^, providing sufficient discriminatory power for differentiating individuals (Waits et al., 2001). P(ID)_sib_ was calculated to be less than 0.01 at 7 loci, therefore we excluded any samples successfully genotyped at fewer than 7 loci (n=14). We also allowed identical samples matching at 7 or more loci to differ at 1 locus to account for genotyping error. In total, we successfully genotyped 385 samples representing 162 unique individuals. 124 unique individuals were identified from KNP (73 males, 51 females). 38 unique individuals were identified from BINP (23 males, 14 females, 1 unknown sex).

### Genetic Diversity

Seven mitochondrial haplotypes were detected across the 162 individuals sequenced. Six of these have been previously detected in Uganda, although five haplotypes represent the first detection of that haplotype in the park from which it was sampled (Table S2). One haplotype was not represented in public databases. Two unique mitochondrial haplotypes were detected in BINP, with 13.2% of individuals carrying F-clade and 86.8% carrying S-clade mtDNA. In KNP, seven unique mitochondrial haplotypes were detected, with 19.4% of individuals carrying F-clade and 80.6% carrying S-clade mtDNA. Across both parks, S-clade sequence LL062 was the most frequently detected haplotype (76.3% of individuals). The AMELY gene was successfully amplified and sequenced in 91 of the 96 unique males identified. In BINP, 100% of the successfully sequenced males carried the forest-typical AMELY gene, while 50.7% of males in KNP carried the forest-typical variant and 49.3% the savanna-typical variant. All four possible combinations of uniparental markers were detected in the males of KNP but not BINP (Table 1).

**Table 1.**
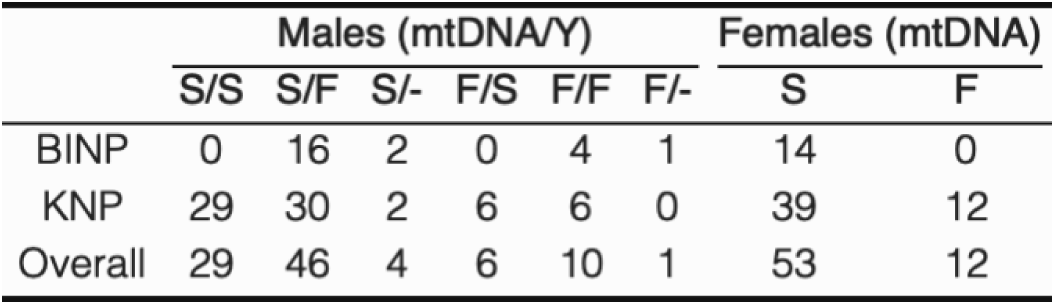
Uniparental sequencing results for the elephants of KNP and BINP. Mitochondrial control region clade assignment and AMELY gene haplotype classification are each presented as “S” (savanna lineage derived) or “F” (forest lineage derived). Males for which the mtDNA control region could be successfully amplified, but the AMELY gene could not, are presented in the “F/-“ and “S/-“ columns.

Genetic diversity values for the 14 microsatellites across KNP and BINP are presented in Table 2. The number of alleles per locus across both parks ranged from 2 at locus Lcy-M27 to 12 at locus FH71. On average, the elephants of KNP had 6.000 ± 0.663 SE unique alleles per locus and of BINP had 4.429 ± 0.429 SE alleles per locus. Two loci in KNP (FH71, LA5) and 1 locus in BINP (LafMS04) deviated significantly from HWE. The inbreeding coefficient (F) was negative for 11 of the 14 loci tested in BINP (mean: -0.095 ± 0.031 SE). This pattern is consistent with a population which has experienced outbreeding. In contrast, only 3 of 14 loci in KNP showed negative F values (mean: 0.049 ± 0.022 SE). KNP and BINP showed distinct clusters along PC1 using Principal Component Analysis (Figure 2), and pairwise F_ST_ between the two parks was 0.11.

**Figure 2.**
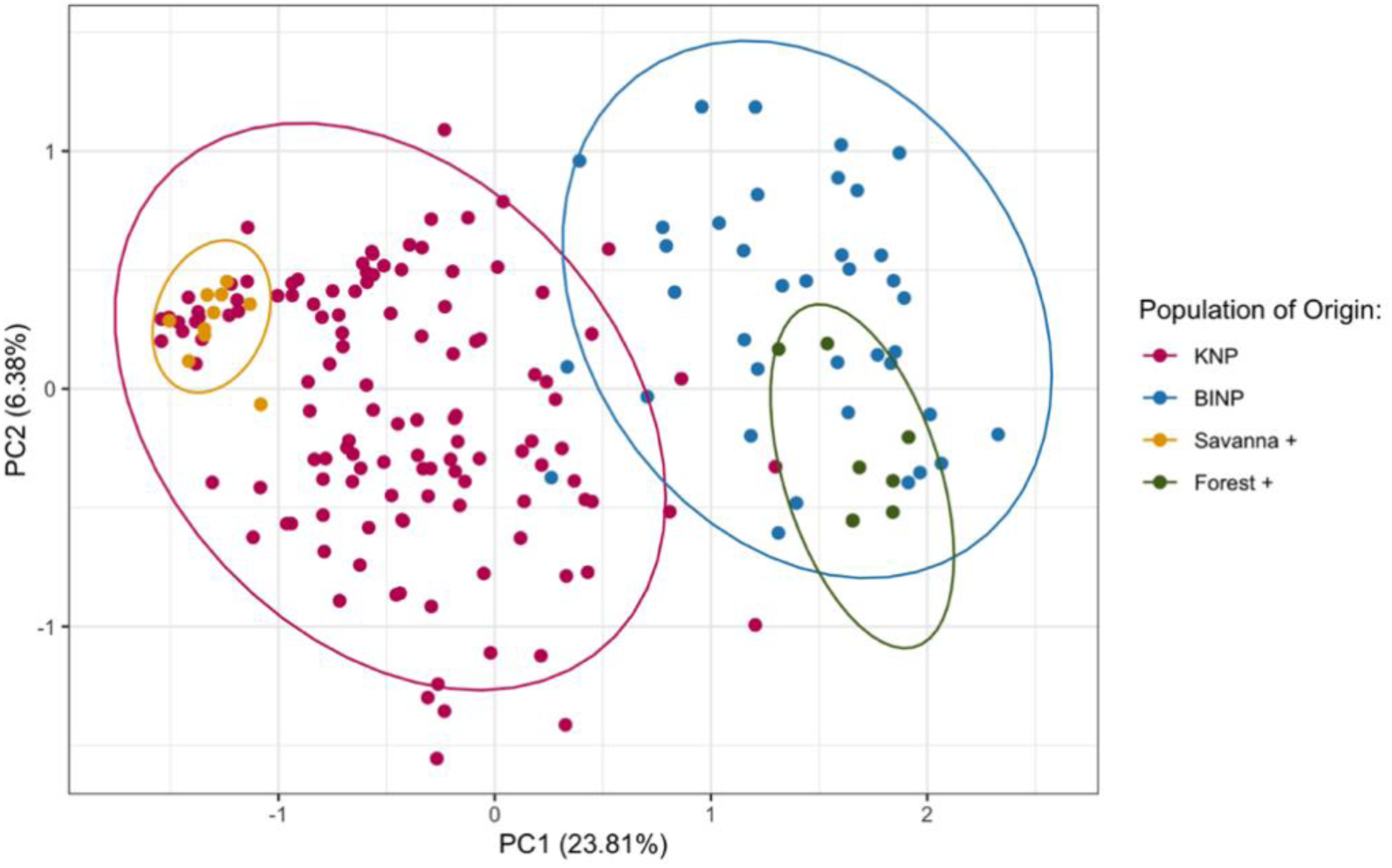
Ordination plot illustrating genetic distance between individuals computed using Principal Component Analysis from 14 microsatellite loci. Dots represent unique individuals, colored by population of origin (KNP, BINP, savanna elephant positive control, forest elephant positive control). Ellipses represent 95% confidence intervals for each group.

**Table 2.**
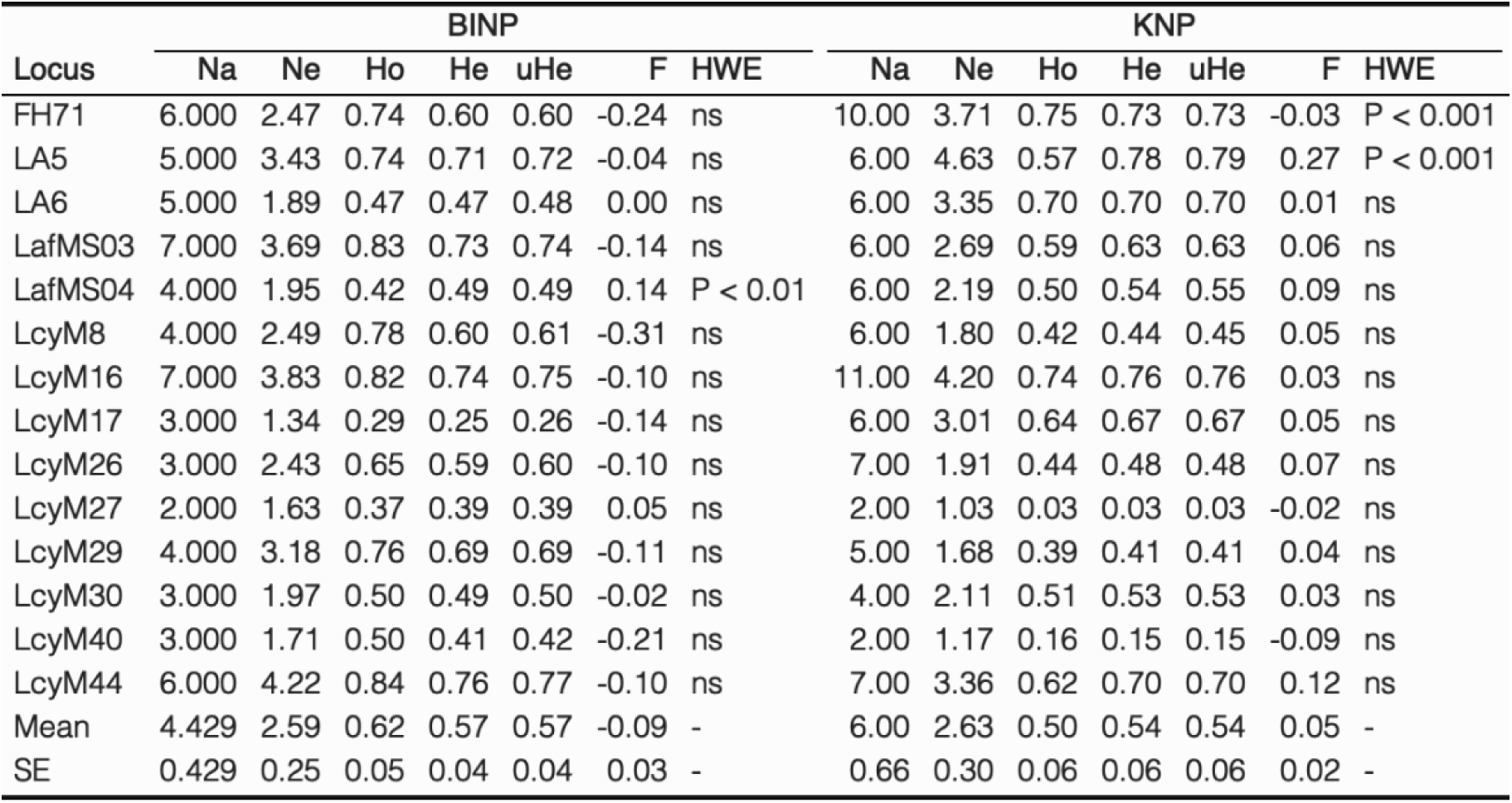
Standard genetic diversity metrics for the elephants of KNP and BINP generated by GenAlEx v. 6.51b2. Abbreviations above are as follows: Na - Number of unique alleles; Ne - Effective number of alleles; Ho - Observed heterozygosity; He - Expected heterozygosity; uHe - Unbiased expected heterozygosity; F - Inbreeding coefficient; HWE - Deviations from Hardy Weinberg Equilibrium.

| Locus | BINP |  |  |  |  |  |  | KNP |  |  |  |  |  |  |
| --- | --- | --- | --- | --- | --- | --- | --- | --- | --- | --- | --- | --- | --- | --- |
|  | Na | Ne | Ho | He | uHe | F | HWE | Na | Ne | Ho | He | uHe | F | HWE |
| FH71 | 6.000 | 2.47 | 0.74 | 0.60 | 0.60 | -0.24 | ns | 10.00 | 3.71 | 0.75 | 0.73 | 0.73 | -0.03 | P < 0.001 |
| LA5 | 5.000 | 3.43 | 0.74 | 0.71 | 0.72 | -0.04 | ns | 6.00 | 4.63 | 0.57 | 0.78 | 0.79 | 0.27 | P < 0.001 |
| LA6 | 5.000 | 1.89 | 0.47 | 0.47 | 0.48 | 0.00 | ns | 6.00 | 3.35 | 0.70 | 0.70 | 0.70 | 0.01 | ns |
| LafMS03 | 7.000 | 3.69 | 0.83 | 0.73 | 0.74 | -0.14 | ns | 6.00 | 2.69 | 0.59 | 0.63 | 0.63 | 0.06 | ns |
| LafMS04 | 4.000 | 1.95 | 0.42 | 0.49 | 0.49 | 0.14 | P < 0.01 | 6.00 | 2.19 | 0.50 | 0.54 | 0.55 | 0.09 | ns |
| LcyM8 | 4.000 | 2.49 | 0.78 | 0.60 | 0.61 | -0.31 | ns | 6.00 | 1.80 | 0.42 | 0.44 | 0.45 | 0.05 | ns |
| LcyM16 | 7.000 | 3.83 | 0.82 | 0.74 | 0.75 | -0.10 | ns | 11.00 | 4.20 | 0.74 | 0.76 | 0.76 | 0.03 | ns |
| LcyM17 | 3.000 | 1.34 | 0.29 | 0.25 | 0.26 | -0.14 | ns | 6.00 | 3.01 | 0.64 | 0.67 | 0.67 | 0.05 | ns |
| LcyM26 | 3.000 | 2.43 | 0.65 | 0.59 | 0.60 | -0.10 | ns | 7.00 | 1.91 | 0.44 | 0.48 | 0.48 | 0.07 | ns |
| LcyM27 | 2.000 | 1.63 | 0.37 | 0.39 | 0.39 | 0.05 | ns | 2.00 | 1.03 | 0.03 | 0.03 | 0.03 | -0.02 | ns |
| LcyM29 | 4.000 | 3.18 | 0.76 | 0.69 | 0.69 | -0.11 | ns | 5.00 | 1.68 | 0.39 | 0.41 | 0.41 | 0.04 | ns |
| LcyM30 | 3.000 | 1.97 | 0.50 | 0.49 | 0.50 | -0.02 | ns | 4.00 | 2.11 | 0.51 | 0.53 | 0.53 | 0.03 | ns |
| LcyM40 | 3.000 | 1.71 | 0.50 | 0.41 | 0.42 | -0.21 | ns | 2.00 | 1.17 | 0.16 | 0.15 | 0.15 | -0.09 | ns |
| LcyM44 | 6.000 | 4.22 | 0.84 | 0.76 | 0.77 | -0.10 | ns | 7.00 | 3.36 | 0.62 | 0.70 | 0.70 | 0.12 | ns |
| Mean | 4.429 | 2.59 | 0.62 | 0.57 | 0.57 | -0.09 | - | 6.00 | 2.63 | 0.50 | 0.54 | 0.54 | 0.05 | - |
| SE | 0.429 | 0.25 | 0.05 | 0.04 | 0.04 | 0.03 | - | 0.66 | 0.30 | 0.06 | 0.06 | 0.06 | 0.02 | - |

### Census Results

We retained 180 unique captures for fecal DNA-based CMR analysis. The population estimate for KNP was 573 individuals (95% CI: 410 to 916). Given the park size of 795-km^2^, crude elephant density was approximately 0.72 elephants/km^2^ (95% CI: 0.52 to 1.03 elephants/km^2^). For BINP, the population estimate was 96 individuals (95% CI: 64 to 145). Crude elephant density for this 331-km^2^ park was 0.29 elephants/km^2^ (95% CI: 0.19 to 0.44 elephants/km^2^).

### Species Identity

All STRUCTURE runs showed the strongest support for two clusters (K=2) with species-specific ancestry (Evanno Method; Figures S1-S2). Because each of our models showed similar trends, we report results from the independent STRUCTURE run without location priors here (see Suppl. for additional models). Pure forest elephant positive controls and pure savanna elephant positive controls showed Q > 0.9 for cluster I and cluster II respectively, while samples collected from the Western Ugandan hybrid zone showed a range of intermediate ancestries (Figure 3). Using a cutoff of Q > 0.9, we classified elephants as pure forest (cluster I), pure savanna (cluster II), or hybrid (intermediate). Of the 124 unique individuals detected in KNP, 22 elephants (17.7%) were classified as pure savanna, 1 elephant (0.8%) as pure forest, and 101 elephants (81.5%) as hybrid. In contrast, in BINP 33 elephants (86.8%) were classified as pure forest, while 5 elephants (13.2%) showed intermediate ancestry. No elephants in BINP were classified as pure savanna elephants.

**Figure 3.**
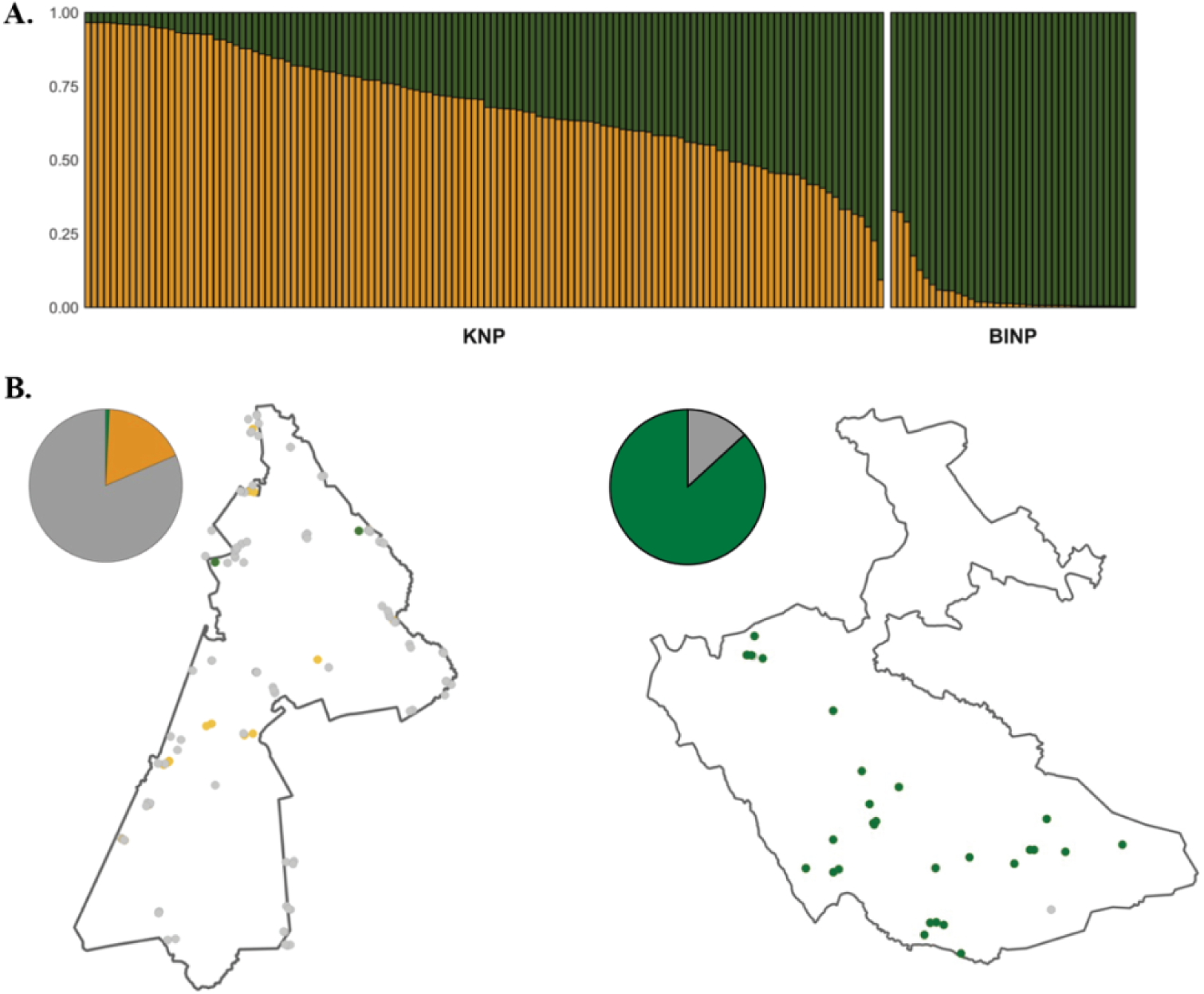
Species classifications for KNP and BINP using 14 microsatellite loci. Green indicates forest ancestry, yellow indicates savanna ancestry, and grey indicates hybrid. A) Plot showing ancestry proportions for the elephants of KNP and BINP inferred separately using the Admixture model in STRUCTURE (K=2) with no location prior. B) Elephant species distribution throughout KNP (left) and BINP (right), colored according to the highest likelihood elephant type (pure forest, pure savanna, hybrid) inferred using Q>0.9.

Next, we used EBHybrids to classify individuals by species and assign each sample a hybrid probability (Table 3). Using the highest stringency likelihood cutoff of 0.95, one elephant (0.8%) in KNP was classified as pure forest, 49.2% as hybrid, 15.3% as pure savanna, and 34.7% could not be assigned to a species bin (“unclassified”). At this same cutoff, the majority of the hybrids in this park were not further classified by hybrid type (86.9%). However, using a more permissive cutoff of 0.5, hybrids were further classified as either savanna backcrosses (85.4%) or F2 hybrids (14.6%). Using a cutoff of 0.95 in BINP, 34.2% of the elephants in BINP were classified as pure forest, 10.5% as hybrids, and 55.3% as unclassified. No pure savanna elephants were detected using this method. Hybrids in BINP were all “unclassified” by hybrid type using a stringency of 0.95 but classified as primarily forest backcrosses (72.7%) using a stringency of 0.5.

**Table 3.** Maximum likelihood species classifications for individuals in KNP and BINP, inferred using EBHybrids (EB: 0.95, 0.8, 0.5) and STRUCTURE (Q>0.9). Hybrid classifications generated by EBHybrids were: F1 hybrid (F1); F2 hybrid (F2); Forest elephant backcross (BXF); Savanna elephant backcross (BXS); Unclassified hybrid (Unc).

| Cutoff | Species |  |  |  | Hybrid Classification |  |  |  |  |
| --- | --- | --- | --- | --- | --- | --- | --- | --- | --- |
|  | Forest | Savanna | Hybrid | Unclassified | F1 | F2 | BXF | BXS | Unc |
| **BINP** |  |  |  |  |  |  |  |  |  |
| EB: 0.95 | 13 | 0 | 4 | 21 | 0 | 0 | 0 | 0 | 4 |
| EB: 0.8 | 22 | 0 | 7 | 9 | 0 | 0 | 3 | 0 | 4 |
| EB: 0.5 | 27 | 0 | 11 | 0 | 3 | 0 | 8 | 0 | 0 |
| Q > 0.9 | 33 | 0 | 5 | 0 | - | - | - | - | - |
| **KNP** |  |  |  |  |  |  |  |  |  |
| EB: 0.95 | 1 | 19 | 61 | 43 | 0 | 2 | 0 | 6 | 53 |
| EB: 0.8 | 1 | 24 | 78 | 21 | 0 | 10 | 0 | 42 | 26 |
| EB: 0.5 | 1 | 34 | 89 | 0 | 0 | 13 | 0 | 76 | 0 |
| Q > 0.9 | 1 | 22 | 101 | 0 | - | - | - | - | - |

## Discussion

### Kibale National Park

The population size and density estimates for KNP generated in this study (573 individuals, 95% CI: 410 to 916; 0.72 elephants/km^2^) are remarkably consistent with the most recent size estimate for this park (566 individuals, 95% CI: 377 – 850; 0.71 elephants/km^2^) which was generated using the traditional LTDS method (Aleper et al., 2021). Given the similar findings between these two methods (see also Hedges et al., 2013; Laguardia et al., 2021a), we argue that fecal DNA-based CMR censuses should be used for potentially admixed populations residing in forests like this one, since this method provides species resolution not captured by LTDS and thus facilitates simultaneous inference of species distribution, hybrid prevalence, and population abundance.

A recent study from the Sebitoli region, a 25-km^2^ area at the northernmost part of KNP, found that 81.3% of the elephants using this region were hybrids and the remaining 18.7% were pure savanna elephants (Bonnald et al., 2021). The absence of any pure forest elephant detections in this previous study was particularly notable given that KNP is a primarily forested park. However, the Sebitoli region is distinctive from other regions of KNP because it is comprised primarily of regenerating forest and constitutes only ∼3% of the park’s total area. For this reason, it remained unclear whether the elephants captured in this region were representative of the park’s elephant population as a whole. Our study expands upon this finding by: 1) extending our genetic survey across the park’s entire area for the first time, and 2) using two uniparentally-inherited markers (mtDNA, AMELY) to infer the sex-linked dynamics of hybridization within KNP. Notably, we identified only one pure forest elephant out of the 124 unique individuals that we sampled (<1%), suggesting that this dearth of forest elephants extends throughout the entire park. As in the Sebitoli region, the majority of the elephants that we detected were also hybrids (81.5%) and savanna elephants (17.7%). These results show that the findings of Bonnald et al. (2021) can be extended to the entire park, which encompasses a variety of different vegetation types. Furthermore, because we detected all four possible combinations of species-typical uniparental markers in the hybrids of KNP and did not detect any F1 hybrids, we suggest that the extensive hybridization characterizing this population happened between pure parental types more than one generation ago, and that hybrids are the offspring of both types of interspecies crosses (female forest x male savanna; female savanna x male forest). These patterns of genetic ancestry are consistent with those previously identified in this hybrid zone (Mondol et al., 2015) and suggest that the hybrid elephants of KNP are fertile and have been mating among themselves and/or backcrossing with their parental taxa for multiple generations, consistent with expectation under a hybrid swarm (Allendorf et al., 2001; Figure S3).

### Bwindi Impenetrable National Park

This study constitutes both the first park-scale, species-level assessment for the elephants of BINP and the first comprehensive elephant census for this park in three decades. The confidence intervals for our CMR model (96 individuals, 95% CI 64 to 145; 0.29 elephants/km^2^) suggest that more elephants are using this landscape than are captured by the current population estimate on record (UWA, 2016). The last systematic survey of the park’s elephants (1992-1993, recce method) put this population at 22 individuals (0.07 elephants/km^2^), including only one adult male, and attributed this small population size to intense poaching throughout the park in the late 1970s (Babaasa, 1994). The Uganda Wildlife Authority’s current population estimate for this park is 43 elephants, estimated from ranger encounter data (UWA, 2016). Hence, it appears likely that there are more elephants in this park than previously thought and that this population has been growing over the last 27 years. Despite differences in census methods over the years, this interpretation is further supported by an annual trend of increased ranger encounter data (UWA, 2016) and an increase in crop raiding by elephants, including at new locations around the park’s boundaries (Butynski, 1986; Natukunda, 2019).

Notably, we also found that the elephants of BINP are primarily forest elephants (86.8%), and the patterns of uniparentally-inherited genetic ancestry suggest a more distant history of hybridization at this site. Although 100% of the detected males at this site carried the expected forest-typical Y chromosome, the population also showed high levels of cytonuclear discordance: 84.8% of putatively pure forest elephants carried savanna lineage derived S-clade mtDNA. Cytonuclear discordance is a well-established phenomenon across the *Loxodonta* genus, but to date this pattern has only been detected in savanna elephant populations carrying forest derived, F-clade mtDNA (Ishida et al., 2011, 2013; Roca et al., 2005). These previous detections are consistent with multiple generations of forest (and subsequent hybrid) females crossing with savanna males and have been explained in terms of the competitive advantage in mating that larger savanna males might have over smaller forest males (Roca et al, 2005). Interestingly, however, we see the opposite pattern in BINP: our findings suggest that the elephants at this site are the result of unidirectional crossing of forest males with savanna (and subsequent hybrid) females. This could be the result of forest males having a context-dependent competitive advantage over savanna males in forested habitats or the introduction of a savanna elephant matriline that over time proceeded to outcompete all forest elephant matrilines. Moreover, this pattern of cytonuclear discord we see would have taken at least 4 generations to achieve (Lavretsky et al., 2016), which would place hybridization at BINP at least 100 years ago.

### Conservation and Management Implications

Our work demonstrates that site-specific, species-level monitoring and management will be critical to developing conservation policy for regions containing admixed elephant populations. While the elephants of western Uganda collectively show high levels of hybridization (Mondol et al., 2015; Kuhner et al., 2025), at the more granular level, species types are non-randomly distributed across different protected areas. For example, while our results suggest the elephants of BINP should be classified and inventoried as forest elephants, the elephants of KNP are most appropriately classified as a hybrid population.

Additionally, further work is needed to better understand the causes and consequences of African elephant hybridization. Specifically, we need to examine how behavior, ecology, health, and fitness of hybrid elephants compares to their parental species in different contexts, investigate putative prezygotic and postzygotic barriers to hybridization, and identify the underlying factors that facilitate and/or maintain elephant hybridization. To this end, we note that the patchy distribution of hybrids and parental types found across the two surveyed sites can be characteristic of disturbance-mediated hybridization (Grabenstein & Taylor, 2018; Mondol et al., 2015). If this is the case, then the ongoing hybridization at KNP could possibly be the result of more recent events, while hybridization at BINP must be related to events that occurred in the more distant past. More work is needed to examine the age of this hybrid zone as it relates to human activity and/or natural ecosystem change.

At the national scale, we emphasize that Uganda is a range state for forest elephants, savanna elephants, and hybrids and should reflect this status as it develops the next iteration of the country’s national elephant conservation management plan, which is set to expire in 2026 (UWA, 2016). Importantly, while our work generates critical data for Ugandan management of KNP and BINP elephants and the classification of these elephants in continent-wide inventories, there are additional forested protected areas throughout Uganda (UWA, 2016) and other parts of Sub-Saharan Africa that face a time-sensitive demand for similar data. In contrast to the LTDS method, our fecal DNA-based CMR method could be smoothly integrated into pre-existing, regularly conducted park surveys by rangers on the ground, enabling census counts and species distributions for elephants throughout protected areas. Such work could also aid in better understanding whether differences in elephant densities between BINP and KNP are due to species identity, population history including poaching, habitat availability, and/or the socioeconomic conditions of surrounding human communities (e.g., De Boer et al., 2013). The increasing elephant densities in the closed forests of KNP (UWA 2016) are particularly concerning given the majority hybrid and savanna elephant species identities, as savanna elephants are known to be ecosystem engineers that create and maintain open vegetation types, although there is no evidence to date that these elephants are driving changes in forest composition (Omeja et al., 2014).

Elephant populations continue to decline across much of the African continent, and a changing climate, poaching, and habitat conversion will likely change the way that African forest and savanna elephants interact and potentially interbreed in locations where their ranges overlap around the Congo Basin (Brandt et al., 2012; Dejene et al., 2021). For these reasons, continued efforts to monitor and understand the effects of hybridization on the elephants in the unique hybrid population of western Uganda and elsewhere will remain critical to continent-wide elephant conservation.

## Author contributions

Study design: CG, NT, DC, CW; Field work coordination, sample collection: CG, DC, JH, RM, MW, EU, CK; Genetic lab work: CG, SB; Data analysis: CG; Writing: CG, DC, CW, DB, PO, CT, MW, CC, RM, JH, NT.

## Acknowledgements

We would like to thank the Ugandan Wildlife Authority and the Ugandan National Council for Science and Technology for granting us permission to conduct this research. Additionally, this work would not be possible without the support of many UWA rangers and field assistants, including: Bonny Balyeganira, Isaiah Mwesige, Jimmy Ogwang, Naume Kiiza, Swaibu Katusabe, Chucknoris Mutegeki and the Greater Virunga Transboundary Collaboration’s 2018 Bwindi-Sarambwe gorilla census team members. We are grateful to the Oregon Zoo for contributing to preliminary work to optimize our sample collection methods for this study. We also thank the San Diego Frozen Zoo for providing African savanna elephant DNA for our use. This work was supported by the National Geographic Society (NGS-388R-18; NGS-90330R-21) and the Morris Animal Foundation (D21ZO-814).

## Ethical standards

This work was conducted under IACUC protocol AUP-21-17.

