## Supplemental Data for "Elephants inhabiting two forested sites in western Uganda exhibit contrasting patterns of species identity, density, and history of hybridization"

**Supplemental Methods**

*Sampling Scheme*

Dung samples were non-invasively collected from across KNP from November 2020 to August 2021, except in February 2021 due to changes in standard operating procedures associated with COVID-19. Sample collection occurred for 6 to 7 days consecutively, every other week. During this time, the field team (3-5 individuals) would set up a base camp out of Uganda Wildlife Authority (UWA) outposts, or when not available, would establish base camps in the forest. The team used informants, crop raiding events, known elephant hotspots, and elephant tracks to search for dung. For each fresh sample, the outside of a minimum of two dung boli were swabbed using an Isohelix Buccal swab, and each swab was placed into a vial containing 1.5 mL of RNAlater (hereafter, “swab samples”). All samples were stored in a cool, dry area in the field for up to 1 week before being transferred to a −20°C freezer.

In BINP, sample collection was implemented over two sampling bouts. The first bout was conducted by the Greater Virunga Transboundary Collaboration from October – December, 2018 as part of a park-wide sweep survey for mountain gorillas and other large mammals (Hickey et al., 2019). Briefly, two field teams (4-5 individuals each) simultaneously surveyed 40 pre-determined units of the park in 14-day shifts by walking reconnaissance routes spaced ~500 meters apart in each unit. When fresh elephant dung piles judged to be <48 hours old were encountered, dung samples were georeferenced and preserved in 15 mL of RNAlater at a 1:1 ratio (hereafter, “pinch samples”). The second sampling bout was conducted by one field team and took place July – September 2019. Sample collection occurred for 5 to 6 days consecutively, every week. After each week, the team would rotate to a different basecamp in a different area of the park. Prior to the survey, a grid of 2x2 km cells was overlaid over a map of the park, and cells containing potential “hotspots” were identified based on vegetation (fruiting trees, bamboo), terrain type (swamps, valleys, water sources) and ranger knowledge of the area. On each survey day, 1-2 non-adjacent hotspot cells were visited and searched by a field team of 2-3 individuals. Cells were not re-visited. Dung piles judged to be <48 hours old were georeferenced and swab samples were taken. All samples were stored in a cool, dry area in the field for up to 6 weeks before being transferred to a −20°C freezer.

*Genetic analyses: Mitochondrial control region and ameloglobin gene*

A 319 bp region of the mitochondrial control region (Brandt et al., 2012) was amplified for all individuals in 25 uL reaction volumes consisting of 12.5 uL Green GoTaq 2X Mastermix, 0.33 uM primers CR-F1 and CR-R2, 0.5 uL BSA, 7.34 uL H2O, and 3 uL DNA. Amplification conditions for this reaction consisted of an initial 3-minute denaturation at 95°C, followed by 35 cycles of 95°C for 30 seconds, 50°C for 45 seconds, and 72°C for 30 seconds, and a final extension step of 72°C for 5 minutes. A 719 bp region of the Y-linked, intronic ameloglobin gene was additionally amplified in two separate reactions for all individuals which were molecularly sexed as males. Each reaction consisted of 10 uL Qiagen Multiplex Mastermix, 6 uL $H_2O$, 0.2 uL BSA, 0.4 uL each of 10X forward and reverse primers (Reaction 1: AmelY-F1/AmelY-R1; Reaction 2: AmelY-F2/AmelY-R2; Mondol, 2015), and 3 uL of DNA. Amplification conditions consisted of an initial denaturation at 94°C for 15 minutes, followed by 40 cycles of 94°C denaturation for 30 seconds, 58°C annealing for 45 seconds, and 72°C extension for 45 seconds, and one final extension step of 72°C for 30 minutes. For each uniparental marker, amplification success was confirmed on a 2% agarose gel, and PCR product which showed a strong band at the expected size range was subjected to an enzymatic cleanup using ExoSAP-IT (Thermofisher) and submitted to Genewiz for Sanger sequencing.

*Genetic Analyses: High-Throughput Amplicon Sequencing*

All samples were genotyped at 14 biparentally inherited microsatellite loci previously shown to be polymorphic in African elephants using a two-step, high-throughput amplicon sequencing (HTAS) approach (Barbian et al., 2018). These loci were: FH71 (Comstock et al., 2000), LA5, LA6 (Eggert et al., 2000), LafMS03, LafMS04 (Nyakaana et al., 2005), and Lcy-M8, Lcy-M16, Lcy-M17, Lcy-M26, Lcy-M27, Lcy-M29, Lcy-M30, Lcy-M40, and Lcy-M44 (Gugala et al., 2016). Primers for microsatellites were ordered with an Illumina sequencing tag attached to the 5' end of both forward and reverse primers (FW primer overhang: 5′ TCGTCGGCAGCGTCAGATGTGTATAAGAGACAG 3′; RV primer overhang: 5′ GTCTCGTGGGCTCGGAGATGTGTATAAGAGACAG 3′).

The 14 loci were then amplified in 2 pooled reactions per sample (Pool 1: LafMS03, LafMS04, Lcy-M8, Lcy-M26, Lcy-M27, Lcy-M29, Lcy-M44; Pool 2: LA5, LA6, FH71, Lcy-M16, Lcy-M17, Lcy-M30, Lcy-M40). DNA was amplified in 15 uL reactions consisting of 7.5 uL Qiagen Multiplex Mastermix, 4.33 uL $H_2O$, 1.2 uL 10X primer pool, and 2 uL DNA. PCRs were performed in a Nexus Gradient thermocycler. Amplification conditions consisted of an initial denaturation step at 95°C for 15 minutes, followed by 45 cycles of 94°C for 30 seconds, 58°C for 90 seconds, and 72°C for 90 seconds. This was followed by a final extension step at 72°C for 10 minutes. After amplification, 5 uL of each of the two pools was combined by sample into 10 uL reactions, subjected to a 1.8X cleanup following the MagBind RxNPure Plus manufacturer's protocol, and diluted 1:100 into $H_2O$.

A second PCR was then performed to ligate dual-indexed adapters to the amplicons. This barcoding PCR was performed in 10 uL volumes consisting of 4 uL Qiagen Multiplex Mastermix, 0.5 uM i5 and i7 adaptors (Nextera XT Index Kit v2; Illumina), and 5 uL diluted amplicon pool (Lepais et al., 2020). Amplification conditions were: 95°C for 5 minutes followed by 10 cycles of 95°C for 30 seconds, 59°C for 90 seconds, and 72°C for 30 seconds. This was followed by a final extension step at 68°C for 10 minutes. Barcoded libraries were cleaned up for a second time at 1.8X following the MagBind RxNPure Plus manufacturer's protocol and quantified using the Quant-IT dsDNA HS kit (Thermofisher). 384 unique dual-indexed libraries were pooled equimolarly per run and sequenced on an Illumina MiSeq at the University of Oregon Genomics & Cell Characterization Core Facility (GC3F) using v3 chemistry at 375 forward read cycles (Barbian et al., 2018). Illumina reads were demultiplexed by sample by the GC3F and adapters were trimmed and filtered for quality (Q>20) using cutadapt v. 3.5 (Martin, 2011). Samples were then demultiplexed by locus and genotype calls were implemented using the program CHIIMP (Barbian et al., 2018). We used a length buffer of 100 bp for locus demultiplexing, a minimum read depth of 150 reads per locus (Salado et al., 2021) and 10 reads per allele for genotype calling. To validate genotype calls, we generated histograms of read lengths for each locus using a custom python script and manually called genotypes based on size. We required that consensus genotypes be confirmed in two or more replicates for heterozygous calls, and in three replicates for homozygous calls.

**Supplemental Data**

| Sample ID | Species | Source |
|---|---|---|
| Lcyc01 | *L. cyclotis* | Gamba Complex, Gabon |
| Lcyc02 | *L. cyclotis* | Gamba Complex, Gabon |
| Lcyc03 | *L. cyclotis* | Gamba Complex, Gabon |
| Lcyc04 | *L. cyclotis* | Gamba Complex, Gabon |
| Lcyc05 | *L. cyclotis* | Gamba Complex, Gabon |
| Lcyc06 | *L. cyclotis* | Gamba Complex, Gabon |
| Lcyc07 | *L. cyclotis* | Gamba Complex, Gabon |
| Lcyc08 | *L. cyclotis* | Gamba Complex, Gabon |
| Lcyc09 | *L. cyclotis* | Gamba Complex, Gabon |
| Lafr01 (SB208) | *L. africana* | Born: San Diego Wild Animal Park<br>Dam: Zimbabwe; Sire: Zambia |
| Lafr02 (SB214) | *L. africana* | Born: San Diego Wild Animal Park<br>Dam: Zimbabwe; Sire: Zambia |
| Lafr03 (SB76) | *L. africana* | Rhodes National Park, Zimbabwe |
| Lafr04 (SB114) | *L. africana* | Uganda (park unknown) |
| Lafr05 (SB527) | *L. africana* | Kruger National Park, South Africa |
| Lafr06 (SB533) | *L. africana* | Kruger National Park, South Africa |
| Lafr07 (SB532) | *L. africana* | Kruger National Park, South Africa |
| Lafr08 (SB528) | *L. africana* | Kruger National Park, South Africa |
| Lafr09 (SB531) | *L. africana* | Kruger National Park, South Africa |
| Lafr12 (SB540) | *L. africana* | Born: San Diego Wild Animal Park<br>Dam: South Africa; Sire: Unknown |

**Table S1.** Origins for positive control DNA used for species assignment analysis in this study. *L. cyclotis* samples (n=9) were collected across the Gamba Complex of Protected Areas in Gabon. *L. africana* samples (n=10) were provided by the San Diego Frozen Zoo (BRG#2020036).

| Haplotype | Clade | Detections: Uganda (Protected Areas) | Detections: Continent-Wide |
|---|---|---|---|
| **LL027** | **F-clade** | **Kibale NP** ** <br> Kidepo Valley NP [1] <br> Murchison Falls NP [1] <br> Queen Elizabeth NP [2] <br> Semuliki NP [2] | DRC [3] <br> Kenya [1] <br> Tanzania [5] |
| **LL062** | **S-clade** | **Bwindi Impenetrable NP** *,2 <br> **Kibale NP** *,2 <br> Queen Elizabeth NP [1,2] | Botswana [5] <br> Eritrea [6] <br> Kenya [1,5,9] <br> Namibia [5,7] <br> Rwanda [2] <br> South Africa [4,5] <br> Tanzania [5] <br> Zimbabwe [5,7,8] |
| **LL068** | **F-clade** | **Kibale NP** ** <br> Queen Elizabeth NP [1,2] <br> Semuliki NP [2] | DRC [2,5] <br> Kenya [4] <br> Rwanda [2] <br> Zambia [5] |
| **LL069** | **S-clade** | **Kibale NP** ** <br> Kidepo Valley NP [1] <br> Murchison Falls NP [1,2] <br> Queen Elizabeth NP [1] | Cameroon [5] <br> Kenya [1,5] |
| **LL094** | **F-clade** | **Bwindi Impenetrable NP** *,2 <br> **Kibale NP** ** | Rwanda [2] |
| **LL099** | **F-clade** | **Kibale NP** ** <br> Semuliki NP [2] | *No detections* |
| **CG001** | **F-clade** | **Kibale NP** ** | *No detections* |

**Table S2.** Geographic distribution data for the mitochondrial haplotypes detected in this study. Seven mitochondrial haplotypes were detected across 156 individuals. Six haplotypes have been previously detected in Uganda, although five haplotypes detected in this study represent the first detection of that haplotype in the park from which it was sampled. One haplotype (CG001) is not currently represented in public databases. [1.] Nyakaana et al., 2002; [2.] Mondol et al., 2015; [3.] Debruyne et al., 2003; [4.] Eggert et al., 2002; [5.] Ishida et al, 2013; [6.] Brandt et al, 2014; [7.] Debruyne et al., 2005; [8.] Charif et al., 2005; [9.] Archie et al., 2006 ***This study**; ****This study only, for submission to NCBI and *Loxodonta Localizer* databases.**

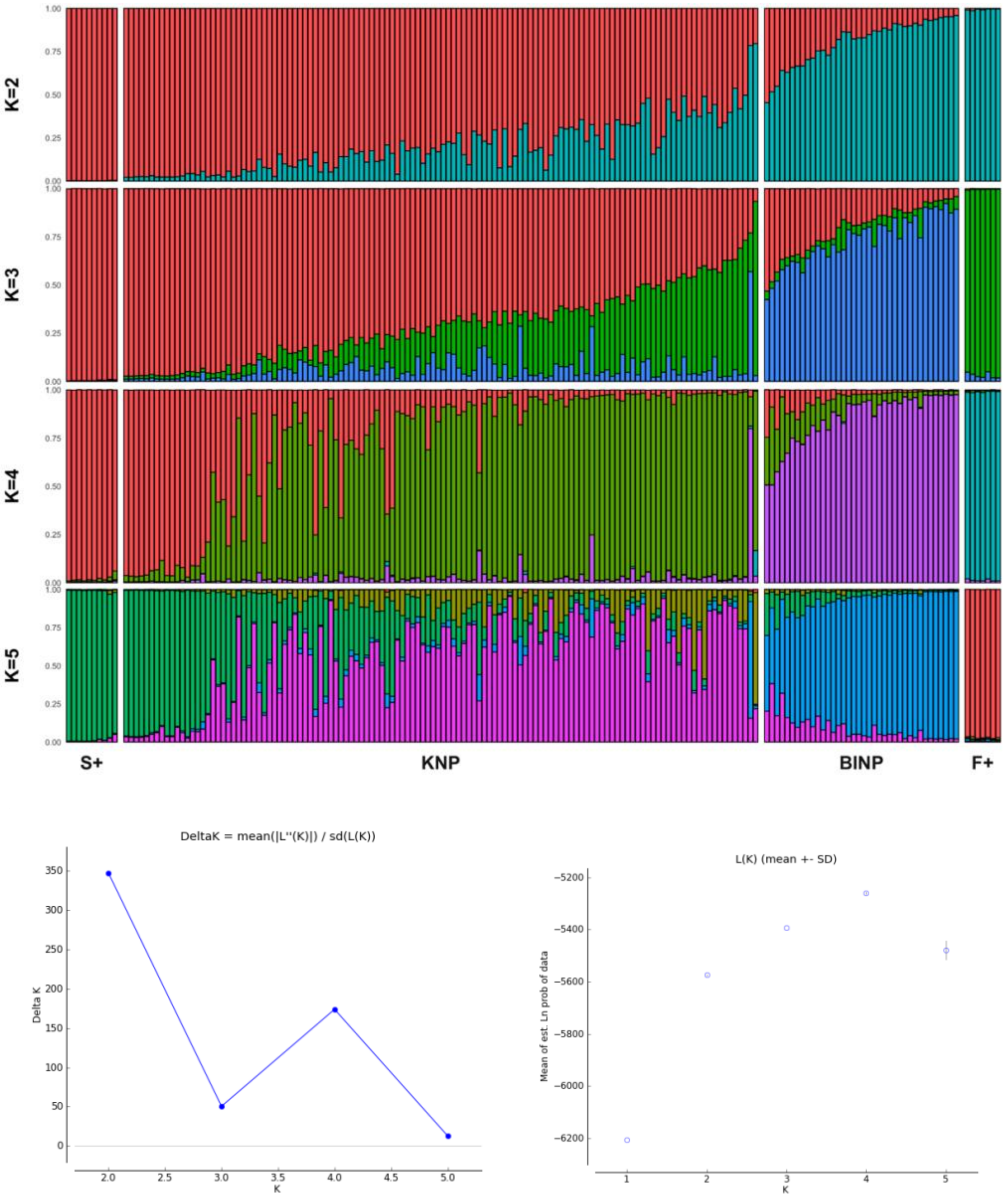


| K | Reps | Mean LnP(K) | Stdev LnP(K) | Ln'(K) | \|Ln''(K)\| | Delta K |
|---|---|---|---|---|---|---|
| 1 | 5 | -6206.040000 | 0.230217 | — | — | — |
| 2 | 5 | -5574.260000 | 1.078425 | 631.780000 | 451.220000 | 418.406559 |
| 3 | 5 | -5393.700000 | 0.924662 | 180.560000 | 46.240000 | 50.007457 |
| 4 | 5 | -5259.380000 | 1.722498 | 134.320000 | 297.100000 | 172.482039 |
| 5 | 5 | -5422.160000 | 59.876272 | -162.780000 | 361.940000 | 6.044798 |

**Figure S1.** STRUCTURE HARVESTER results for the combined STRUCTURE run implemented with location priors for K=2 through K=5. Implementation of the Evano method indicates maximum support for population structure at K=2. Abbreviations above are: KNP = Kibale National Park; BINP=Bwindi Impenetrable National Park; F+ = Forest elephant positive control; S+ = Savanna elephant positive control.

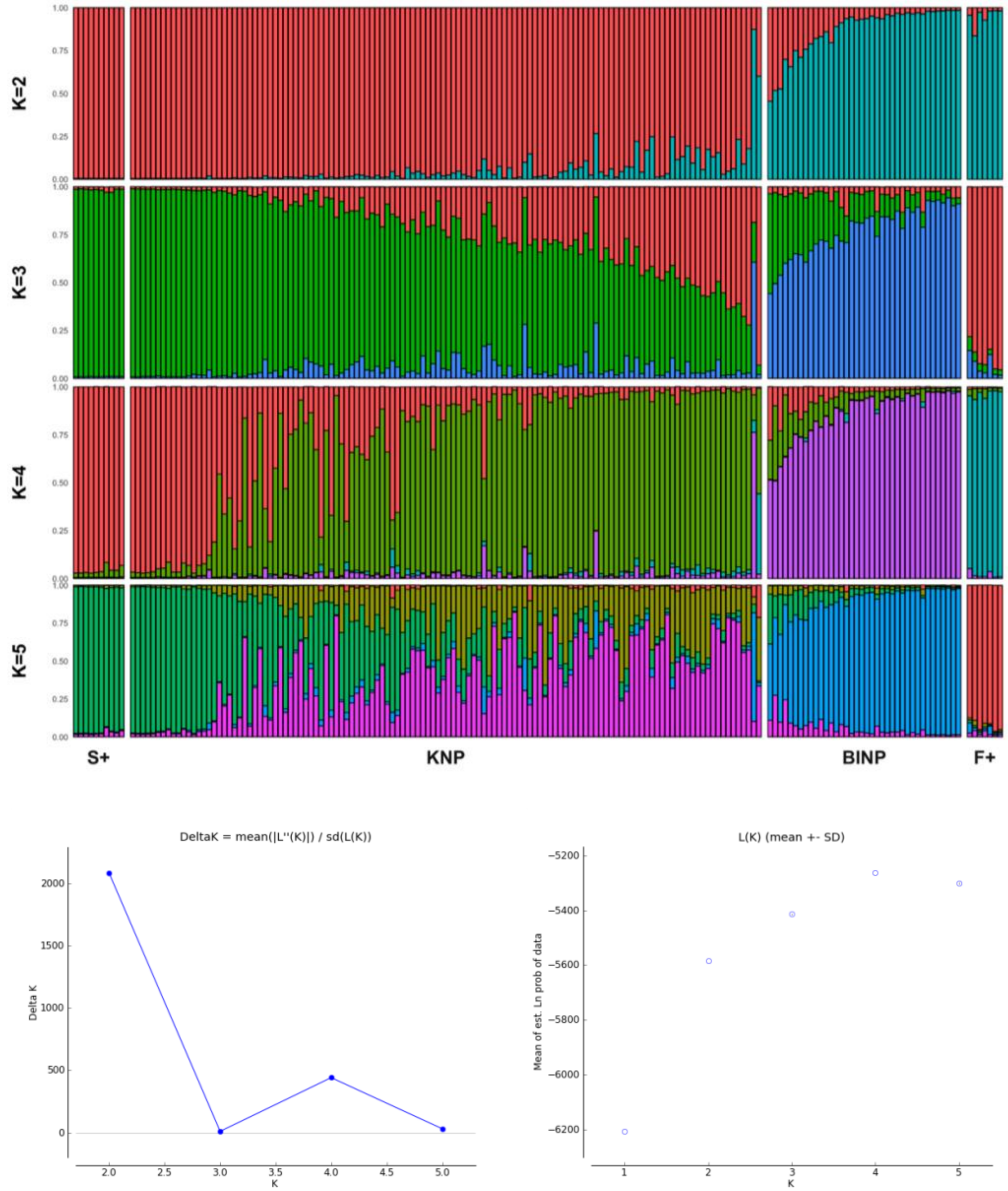


| K | Reps | Mean LnP(K) | Stdev LnP(K) | Ln'(K) | \|Ln''(K)\| | Delta K |
|---|---|---|---|---|---|---|
| 1 | 5 | -6206.060000 | 0.270185 | — | — | — |
| 2 | 5 | -5584.820000 | 0.258844 | 621.240000 | 449.480000 | 1736.492736 |
| 3 | 5 | -5413.060000 | 1.957805 | 171.760000 | 20.960000 | 10.705868 |
| 4 | 5 | -5262.260000 | 0.391152 | 150.800000 | 185.820000 | 475.058114 |
| 5 | 5 | -5297.280000 | 4.399659 | -35.020000 | 146.550000 | 33.309399 |

**Figure S2.** STRUCTURE HARVESTER results for the combined STRUCTURE run implemented without location priors for K=2 through K=5. Implementation of the Evano method indicates maximum support for population structure at K=2. Abbreviations above are: KNP = Kibale National Park; BNP=Bwindi Impenetrable National Park; F+ = Forest elephant positive control; S+ = Savanna elephant positive control.

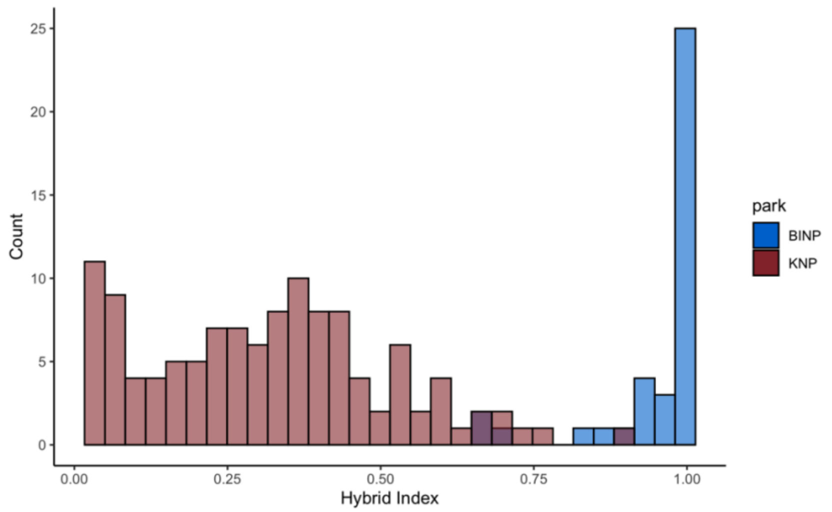


**Figure S3.** Distribution of hybrid indices for the elephants of KNP and BINP. A hybrid index of 0 denotes pure savanna ancestry, and a hybrid index of 1.0 denotes pure forest elephant ancestry. The elephants of KNP show a broad range of intermediate ancestries consistent with expectation under a hybrid swarm (Allendorf et al., 2001), while the elephants of BINP show a narrow, unimodal distribution around forest elephant ancestry.
